# The territory of my body: Testosterone prevents limb cooling in the Rubber Hand Illusion

**DOI:** 10.1101/279463

**Authors:** Donné van der Westhuizen, Teneille Page, Mark Solms, Jack van Honk

## Abstract

The Rubber Hand Illusion (RHI) is an experimental paradigm for assessing changes in body ownership. Recent findings in the field suggest that social-emotional variables can influence such changes. Since body ownership is a mechanism that enables subjects to organise information about the environment and interact efficiently in it, changes in body ownership should have important implications for social power. In the current study, we investigated whether 0.5mg of testosterone, a steroid hormone strongly implicated in the achievement of power and dominance, affected the RHI. Forty-nine females participated in a double-blind, placebo-controlled experiment in which the RHI was induced. Compared to placebo, testosterone had no effects on the usual alteration of subjective ownership over the rubber limb or on subjective sense of proprioceptive drift. However, unlike the placebo group, testosterone-treated participants did not display an objective decline in the temperature of their own (hidden) hand following induction of the illusion. These findings suggest that testosterone strengthens implicit but not explicit bodily self-representations. We propose that effective maintenance of implicit body boundaries can be regarded, conceptually, as a primary defensive state facilitating integrity of the self.

## 1. INTRODUCTION

Several decades of research point to the steroid hormone, testosterone, as a key determinant of socially dominant behaviors that promote self-interests and the acquisition of power and high status in group hierarchies (Carré & Putman, 2010; van der Westhuizen & Solms, 2015a; Ronay & von Hippel, 2010). For example, testosterone facilitates resilience in stressful situations (Enter, Spinhoven & Roelofs, 2014; Hermans et al., 2007; Russo et al., 2012) and increases the motivation to approach social challenge as a means of achieving dominance in potentially hazardous social contexts (Terburg & van Honk, 2013). Some studies indicate that testosterone also regulates certain pro-social forms of behavior too, but the defensive monitoring of implicit social threat (Hermans, Ramsey & van Honk, 2008; Terburg et al., 2012; van Honk et al., 1999) and the attenuation of certain empathic processes that might otherwise interfere with competition (Hermans et al., 2006) have been established as key mechanisms via which it regulates power and social status.

Currently, though the neural chemistries and patterns of behavioral gestures that support the acquisition of social power are well described in both humans and other species (for a review, see van der Westhuizen & Solms, 2015b), research is only now beginning to elucidate the sensory-motor profiles of the body that are modulated by testosterone in order to influence personal power. For instance, a recent study by our group found that testosterone increases the implicit experience of motor control over goal-directed actions (van der Westhuizen et al., 2017). Here, female participants administered 0.5mg of testosterone experienced enhanced ownership over the effects of their actions, a phenomenon referred to as the sense of agency. In other recent work, Moeini-Jazani et al., (2017) showed that increases in social power lead to improved perception of visceral bodily signals, argued by the authors to reflect a shift toward an internal locus of control. Together, these findings suggest that the sensation of power and control may manifest firstly in the bodily domain before extending upwards to influence higher aspects of cognition and feeling.

This idea falls within the increasingly influential embodied cognition framework (Barsalau, 2008; Wilson, 2002), in which the body is viewed as a constituting factor in mental experience. Representations of the body therefore have direct implications for higher-order aspects of self-consciousness and bodily signals of all kinds – whether affective, sensory or homeostatic – have been argued in this way to constrain one’s experience in the world, even when the signals are outside of awareness (Gallagher & Aguda, 2015). Activating the muscles through bodily gestures, like making a fist, for instance, tends to make men feel more powerful (Schubert & Koole, 2009). Because we draw directly on our bodily phenomenology to understand and perceive the world, the brain mechanisms that regulate body ownership should therefore have significant consequences for psychological experiences related to social power, especially during exchanges involving *other* bodies.

### 1.1 Body Ownership

Body ownership, reflected by the feeling that “I am the subject of this thing, my body” is argued to provide the basic foundation for a stable sense of self (Damasio, 2012; Tsakiris, 2016). In this way, bodily ownership is integral to adaptive emotional functioning (James, 1905). At around 18 months of age, children begin to recognise themselves as distinct social entities (Rochat, 2003), imbued with the feeling that this acting body is “me” and those things beyond its confines are “not me”. Though this phenomenon tends to be taken for granted in everyday experience, body ownership is sustained by a complex network of brain systems that represent the external and internal body (Tsakiris, 2016). These representations are dynamic and hierarchical, involving the registration and integration of an extensive range of sensory (visual, somatosensory, auditory, vestibular, visceral), motor and affective signals (Damasio 2000; de Vignemont, 2011; Ehrsson, Spence & Passingham, 2005; Metzinger 2003). In general, the synchronous registration and integration of these bodily signals within insula (Farrer & Frish, 2002; Tsakiris et al., 2007), pre-motor and cerebellar brain regions (Ehrsson, Holmes & Passingham, 2005) has been shown to give rise to the subjective feeling of body ownership (Lopez, Halje & Blanke, 2008; Ehrsson, 2012; Tsakiris, 2008; 2010).

This continuity between the self and the body, in relation to others and the environment, not only establishes self-location from an egocentric visuo-spatial perspective, but it is also a mechanism that allows the subject to organize information and interact efficiently and with a sense of familiarity in rapidly changing settings (Neisser, 1988; Metzinger, 2003). As such, bodily self-consciousness must play an important role in adaptive, “approach” human behavior, especially so in that it provides a reference point from which to appraise and actively respond to different kinds of stimuli in the immediate environment, including those that might be socially threatening.

### 1.2 Malleable body schemas and the RHI

‘Threat’, in fact – to the integrity of the bodily self – has played a key role in the study of changes in body ownership, which has been extensively investigated in a range of procedures that experimentally induce the feeling of ownership over “not me” body parts and even whole bodies (Salomon et al., 2013; Tsakiris, 2016). Several experimental paradigms that manipulate multi-sensory integration have revealed that bodily self-representations are not rigid but that individuals differ in the extent to which their existing schema is malleable due to its multi-sensory nature and can be *replaced* by “not me” bodies and body parts (Botvinick & Cohen 1998; Constantini & Haggard, 2007; Tsakiris and Haggard 2005; Tsakiris et al. 2007).

In an experimental procedure referred to as the rubber hand illusion (RHI; Botvinick & Cohen 1998), synchronous stroking of one’s own unseen hand and a rubber hand that is anatomically aligned with the body, leads to a feeling of ownership over the prosthesis as well as changes in the existing bodily matrix. This is thought to occur because of visual dominance in multisensory integration (Tsakiris and Haggard 2005), an effect that appears to be robust to distinctions in size (Preston & Ehrsson, 2014), age (Banakou, Groten & Slater, 2013), gender (Slater, et al., 2010) and race (Maister, Sebanz, Knoblich & Tsakiris, 2013). The extent of the illusion can be quantified in three ways: by the subjective, self-reported feeling of ownership (Botvinick & Cohen 1998; Longo et al., 2008), by a shift in subjective perception of the real limb’s location towards the rubber hand (proprioceptive drift; Botvinick & Cohen 1998) and by an objective drop in temperature in the real hand (Hohwy & Paton, 2010; Kammers, Rose & Haggard, 2011; Moseley et al., 2008; Thakkar, Nichols, McIntosh & Park, 2011; Tsakiris, Tajadura-Jiménez & Constantini, 2011; Van Stralen et al., 2014).

Moseley et al (2008) have suggested that a drop in limb temperature during the RHI can be interpreted as a disruption in the physiological regulation of the real hand, suggesting that homeostatic processes are affected by subjective ownership over the rubber (versus the real) hand. This reflects, on an implicit level, a disturbance in conception of the bodily self. Indeed, reductions in temperature have also been observed in the full-body illusion (Salomon et al., 2013). In support of Moseley’s et al. (2008) view, several studies have shown that motor and sensory nerve conductivity and responsivity declines in response to cooling (Abramson et al., 1966; Dioszeghy & Stalberg, 1992; Halar, DeLisa & Soine, 1983; Herrera et al., 2010). In fact, dell Gatta et al., (2016) report significantly reduced motor evoked potentials in the real hand following induction of the illusion and Barnsley et al. (2011) have demonstrated that histamine reactivity in the real hand of participants increases in response to the illusion. This implies that not only is ownership over a foreign limb attained, but that the motor and interoceptive systems react by “disowning” the real hand, as if it had become a foreign body (Tsakiris, 2017).

It is not yet clear whether these changes in minimal bodily consciousness have any meaningful bearing on adaptive, goal-oriented behaviour. Evidence does, however, suggest that blurring of self-other boundaries appears to be a function of socio-emotional variables, with several studies showing that ‘concern for others’, or empathic motivation, positively predict degrees of malleability in the subjective domain (Asai et al., 2011; Cascio et al., 2012; Paton et al., 2012). For instance, Ide and Wada (2017) recently found that salivary oxytocin, which is well known for its role in social nurturance and empathy (Panksepp, 1998), positively relates to subjective ownership in the RHI. Testosterone may therefore disrupt bodily plasticity, given its interference with empathic processing (Hermans et al., 2006).

From a social perspective, a flexible conception of the self may facilitate protective behaviours away from the singular-self and toward another individual via the merging of the body’s own “cues” for defence with representations of the other (Ehrsson et al., 2007). Individuals with malleable body representations may in turn have difficulty prioritizing their own needs. A study by Paladino et al. (2013) on multisensory integration in the enfacement illusion, a facial analogue of the RHI, showed that following the induction of an illusory feeling of ownership over another person’s face, participants were more likely to conform and defer to the judgement of this embodied “other”. People who lack experiences of power in their daily living also demonstrate lower interoceptive accuracy (Moeini-Jazani et al., 2017) – a predictor of more malleable body schemas (Tajadura-Jiminez et al., 2011) which is also closely linked to psychiatric disturbance (Bender et al., 2007; de Bonis et al., 1995; Noel et al., 2017). This may occur because disembodiment reduces the flow of relevant bodily information that a subject uses to monitor how well they are faring in a current situation (Pezzulo, Rigoli & Friston, 2015).

In keeping with embodied cognition theories, disowning the body thus appears to relate in fundamental ways to emotional functioning and the fact that not all RHI studies (David et al., 2014; Dehaan et al. 2017; Grynberg & Pollatos, 2015; Paton et al., 2012; Rohde et al., 2013; Thakkar et al., 2011) report drops in limb temperature further supports a role for specific emotional variables as contingencies in this regard. If such variables are not controlled for, they may cancel out at the group level any reliable change in homeostatic regulation. Thus, the link between social emotions and body ownership strongly suggests a role for testosterone in the modulation of corporeal experience.

### 1.4 Overview and aims

In sum, the RHI is well established as an experimental paradigm for undermining the integrity of the bodily self. The body has direct implications for mental experience because we draw on bodily states to evaluate and respond to “not me” objects in the external environment. The capacity for embodiment is therefore potentially empowering and disembodiment, by contrast, could be argued to reflect a state of disempowerment. Here we explore the idea that effective maintenance of the body’s representational boundaries can be regarded, conceptually, as a defensive state facilitating integrity of the self. Using a double-blind, placebo-controlled design, this study examined the effects of 0.5mg testosterone on three indices of the RHI in a sample of women, including subjective feelings of ownership, proprioceptive drift and objective changes in limb temperature.

## 2. METHODS

Ethical approval was granted by the University of Cape Town’s Human Research Ethics committee (HREC REF 868/2014). All data was collected in accordance with the Declaration of Helsinki. Informed consent was obtained prior to commencement of the study and a debriefing took place upon completion of data collection. There were no reports of negative side effects from the testosterone or placebo administration and no participant withdrew from the study.

### 2.1 Participants

49 neurologically healthy, right-handed female participants, aged between 18 and 30 years were recruited to participate in the study in exchange for 18 USD. Testing was performed during the pre-ovulatory phase of the menstrual cycle since androgen levels are relatively constant during this time. Individuals who were taking any form of hormonal contraception or other form of medication were excluded from participation, as were those with a history of psychiatric illness or with a BMI over 30. We excluded males as the reliability of the testosterone administration protocol has only been established in females (Tuiten et al., 2000).

### 2.2 Self-report measures

#### Profile of Mood States (POMS) abbreviated questionnaire

We employed the POMS (Shacham, 1983) to assess whether indices of the RHI in the placebo group were influenced by current mood state, since positive emotions may be a proxy for social coping and resilience. The POMS consists of 40 adjectives that measure depression, anger, confusion, vigour and fatigue. It was completed at the beginning of the experimental session. A Total Mood Disturbance (TMD) score was calculated by adding together the scores for Tension, Depression, Anger, Fatigue and Confusion and then subtracting the score for Vigour.

#### Self-report questionnaire for the RHI

Following Tsakiris, Tajadura Jiménez and Costantini (2011), who abridged and validated Longo’s (2008) original 27-item self-report questionnaire for the RHI, we used their 8-item version. This self-report questionnaire for the RHI provides a measure of the subjective experience of the induction of the illusion and was based on a qualitative study where five participants were encouraged to freely report their experiences after induction. The abridged version focusses on themes relating specifically to body ownership (first 5 items) and location of the real hand in relation to the rubber hand (last 3 items), and makes use of a Likert scale where scores can range from 1 (strongly disagree) to 7 (strongly agree) with 4 being neither agree nor disagree, amounting to a total possible score of Refer to Table 1 for a list of these 8 items.

**Table 1:** Items from the RHI self-report questionnaire.

|  |  |
| --- | --- |
| 1 | It seemed like I was looking at my own hand, rather than at a rubber hand. |
| 2 | It seemed like the rubber hand was a part of my body. |
| 3 | It seemed like the rubber hand was my hand. |
| 4 | It seemed like the rubber hand belonged to me. |
| 5 | It seemed like the rubber hand began to resemble my real hand. |
| 6 | It seemed like the rubber hand was in the location where my hand was. |
| 7 | It seemed like the touch I felt was caused by the paintbrush touching the rubber hand. |
| 8 | It seemed like my hand was in the location where the rubber hand was. |

Items 1-5 gauge subjective ownership more explicitly and can be scored separately out of 35.

### 2.3 Apparatus and procedures

Participants were randomly assigned to either the testosterone or placebo group and tested on one occasion in a double-blind, between-subjects design. To control for hormonal fluctuations in diurnal cycles, testing sessions were standardised to the afternoon. Participants reported to the lab exactly 4 hours prior to the experimental session at which time they received sublingual administration of testosterone or placebo. This schedule was based on previous research which has shown the efficacy of sublingual testosterone administration to peak after 4-6 hours (Hermans et al., 2006; 2007; 2008; Montoya et al., 2013; Terburg et al., 2012; Tuiten et al., 2000; van Honk, 2004; 2005). The testosterone administration consisted of 0.5mg of testosterone, 5mg of the carrier hydroxypropyl beta cyclodextrin, 5mg ethanol and 5ml of water. For the placebo administration, the 0.5mg of testosterone was omitted. The administrations were identical in taste and delivered blind in coded vials. During the interval, participants were requested to refrain from engaging in strenuous or sexual activity, to avoid smoking or consuming excessive caffeine or consuming a heavy meal less than one hour prior to the experiment.

The RHI was carried out in accordance with standard procedure (Tsakiris & Haggard, 2005) and operationalised in terms of a self-report questionnaire, measures of proprioceptive drift and temperature change in the experimental limb. The researcher and participant sat opposite one another at a table, separated by a rectangular box with openings on either side so that the researcher could see the participant’s left hand resting inside the structure (See Figure 1). The rim of the top surface of the box on the side facing the experimenter included a ruler to track hand position estimates. A life-like, plastic, left-hand prosthesis was placed next to the box in the same orientation as the participant’s real left hand at a distance of approximately 30cm away. Prior to placement, participants were presented with three prostheses, differing in terms of skin shade, and asked to select for use the one which they felt most closely resembled their own skin colour. A dark smock was draped over the participant’s shoulder to hide the position of their real left arm, leaving only the rubber hand visible.

**Figure 1.**
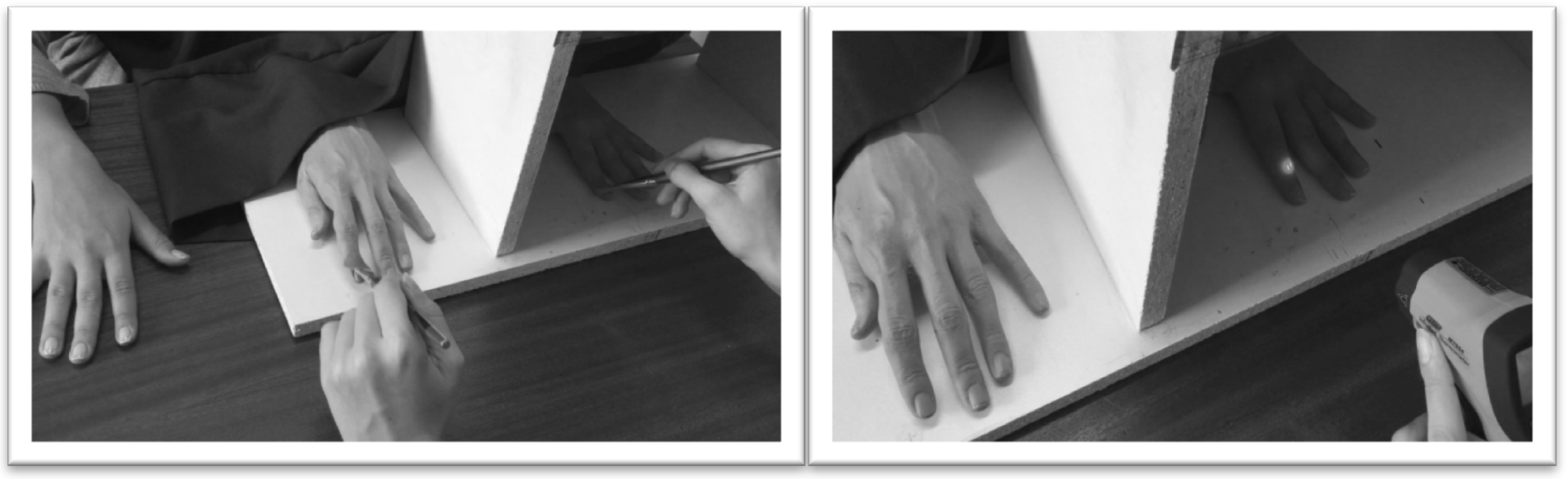
The set up for the RHI requires that the participant is not able to see their real left hand while the prosthesis is in clear sight. For proprioceptive judgements, participants were questioned about the current location of their hand relative to position “1”. Temperature readings by infra-red thermometer were taken from location “2”. The box measured 40cm in width, 28cm in height and 23cm in depth.

The RHI task involved two blocks – an experimental-synchronous condition and a control-asynchronous condition – which were counterbalanced across participants. At the start of each block, the participant was instructed to place their left hand inside the box and look towards the prosthesis. Prior to induction, the RHI questionnaire was administered and a proprioceptive judgment was obtained by asking participants to use their free, right hand to indicate along the top surface of the box where they felt the nail of their left index finger was currently located. This location was quantified in terms of the corresponding location of the ruler, which was only visible to the researcher. A baseline temperature reading of the knuckle closest to the nail of the left hand index finger was then taken using an MT964 non-contact infrared thermometer (Major-Tech Ltd.), which displays precision up to two decimal places. Following Tsakiris, Tajadura-Jiménez and Constantini (2011), only temperature on the experimental limb was recorded.

The first induction phase of the illusion was then performed for 120 seconds, during which the experimenter applied brush strokes at a speed of 1Hz using two identical paintbrushes to the participant’s hidden hand and the prosthesis, to which, their gaze was fixed. Participants were instructed to remain still and refrain from talking during the induction. For the synchronous block, the brush strokes were administered at the same time, in synchrony, running from the knuckle of the index finger to the fingertip. In the asynchronous block, brushstrokes to the prosthesis were applied 180 degrees out of phase (stroking began on the rubber hand between 0.5-1 second before tactile stimulation of the real hand).

Immediately following the stroking phase, a second temperature reading was taken and the participant was requested to make another proprioceptive judgment before removing their hand from the box and filling in the RHI questionnaire once more. Between blocks, participants were asked to briefly stand up so as to encourage the “re-setting” of sensory motor parameters between stroking conditions.

**Figure 2.**
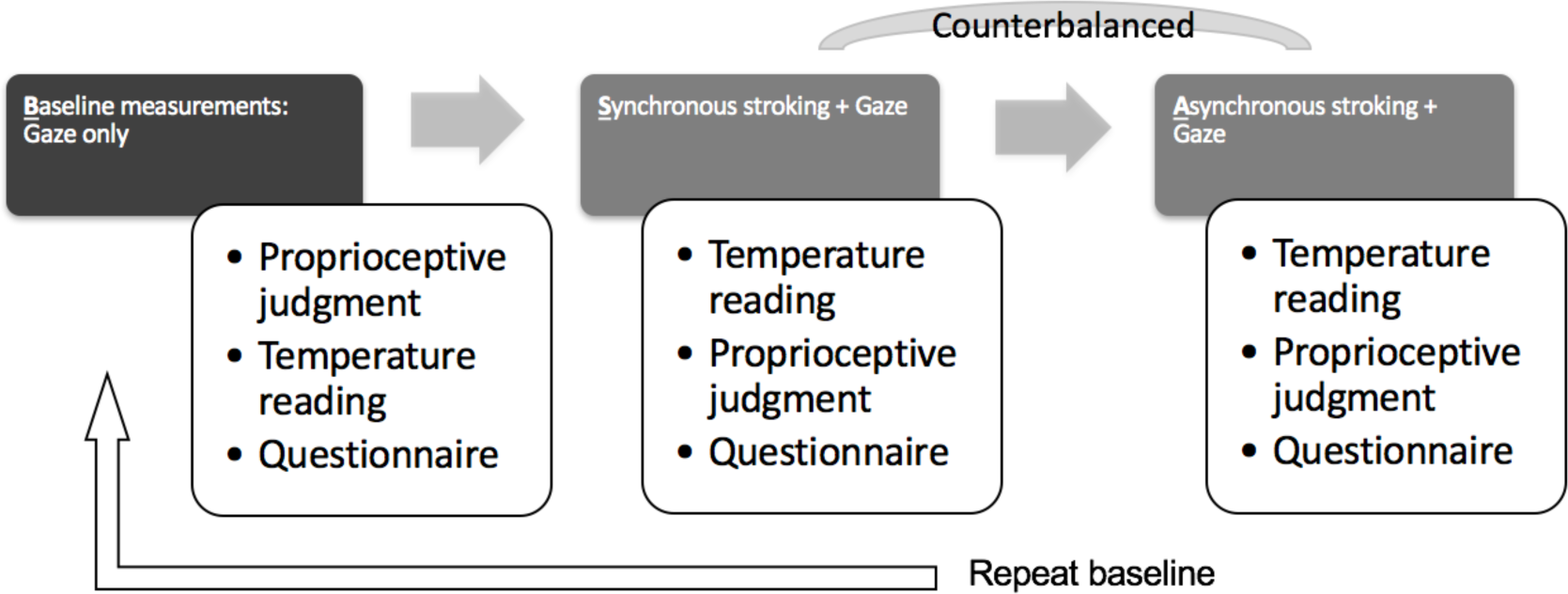
Schematic overview of the procedure and recording schedule for subjective ownership, proprioceptive and physiological temperature changes during the RHI induction. Baseline measures are recorded prior to induction in each stroking condition to control for changes attributed solely to the observation of the prosthesis (visual capture).

### 2.4 Analysis of the RHI

The RHI procedure results in several indices that allow one to control for both visual capture, i.e., the effect on illusory changes occurring as a result of simply processing the visual properties of the prosthesis without simultaneous tactile input, as well as the effect of multisensory integration induced specifically by synchronous stroking. The first baseline reading at the very start of the experiment was used to compare existing differences on each measure across placebo and testosterone groups.

Thus, to compute scores that control for visual capture and which represent the change as a result of stroking in both conditions, baseline measures prior to stroking of the subjective experience of the illusion, proprioceptive judgements and temperature readings weresubtracted from post-induction scores in each stroking condition. For proprioception, this value is referred to in the literature as “proprioceptive drift”. These scores, which we will call ‘baseline corrected change’ (BCC), were then used to calculate the specific influence of multi-sensory integration, which is of key relevance to the manifestation of body ownership. These critical values are referred to as the Perceptual Shift and are calculated by subtracting scores from the Asynchronous condition from scores in the Synchronous condition. Refer to Table 2 for an overview of these differentials.

**Table 2.**
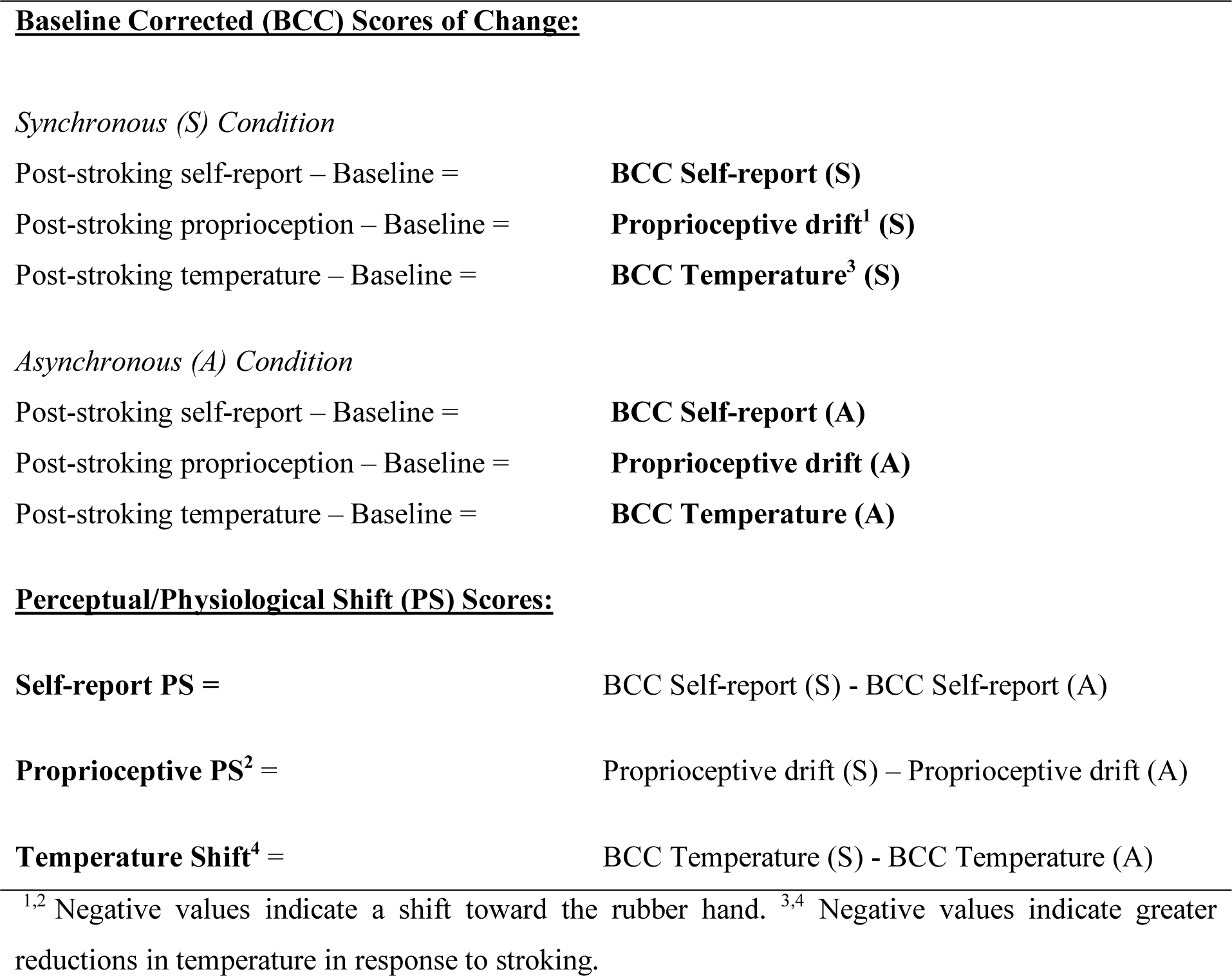
Calculation of differential variables for analysis

|  |  |
| --- | --- |
| Post-stroking self-report – Baseline = | **BCC Self-report (S)** |
| Post-stroking proprioception – Baseline = | **Proprioceptive drift<sup>1</sup> (S)** |
| Post-stroking temperature – Baseline = | **BCC Temperature<sup>3</sup> (S)** |
| *Asynchronous (A) Condition* |  |
| Post-stroking self-report – Baseline = | **BCC Self-report (A)** |
| Post-stroking proprioception – Baseline = | **Proprioceptive drift (A)** |
| Post-stroking temperature – Baseline = | **BCC Temperature (A)** |
| **Perceptual/Physiological Shift (PS) Scores:** |  |
| **Self-report PS =** | BCC Self-report (S) - BCC Self-report (A) |
| **Proprioceptive PS<sup>2</sup> =** | Proprioceptive drift (S) – Proprioceptive drift (A) |
| **Temperature Shift<sup>4</sup> =** | BCC Temperature (S) - BCC Temperature (A) |

<sup>1,2</sup> Negative values indicate a shift toward the rubber hand. <sup>3,4</sup> Negative values indicate greater reductions in temperature in response to stroking.

## 3. RESULTS

Prior to analysis, the data was screened and outliers were removed if they exceeded 3 standard deviations plus/minus the mean. Review of the self-report data revealed 1 outlier in each group. A further outlier was found in the proprioception data. 1 participant failed to provide a full data set for the self-report questionnaire. Data sets from 2 further participants were excluded due to noncompliance with the experimental instructions (movement), leaving a total sample of 46 for temperature and proprioception indices and 44 in the self-report pool.

The testosterone and placebo groups did not differ significantly on any measure of the RHI at baseline (Self-report questionnaire: *t*_42_ = .32, *p* = .75; Proprioceptive mislocalisation: *t*_44_ = 1.59, *p* = .87; Limb temperature: *t*_44_ = .81, *p* = .42. Mean values are displayed in Table 3. This implies that the groups were comparable at the start of the experiment in terms of their hand temperature, their proprioceptive representations and their subjective perceptions of the prosthesis.

**Table 3.**
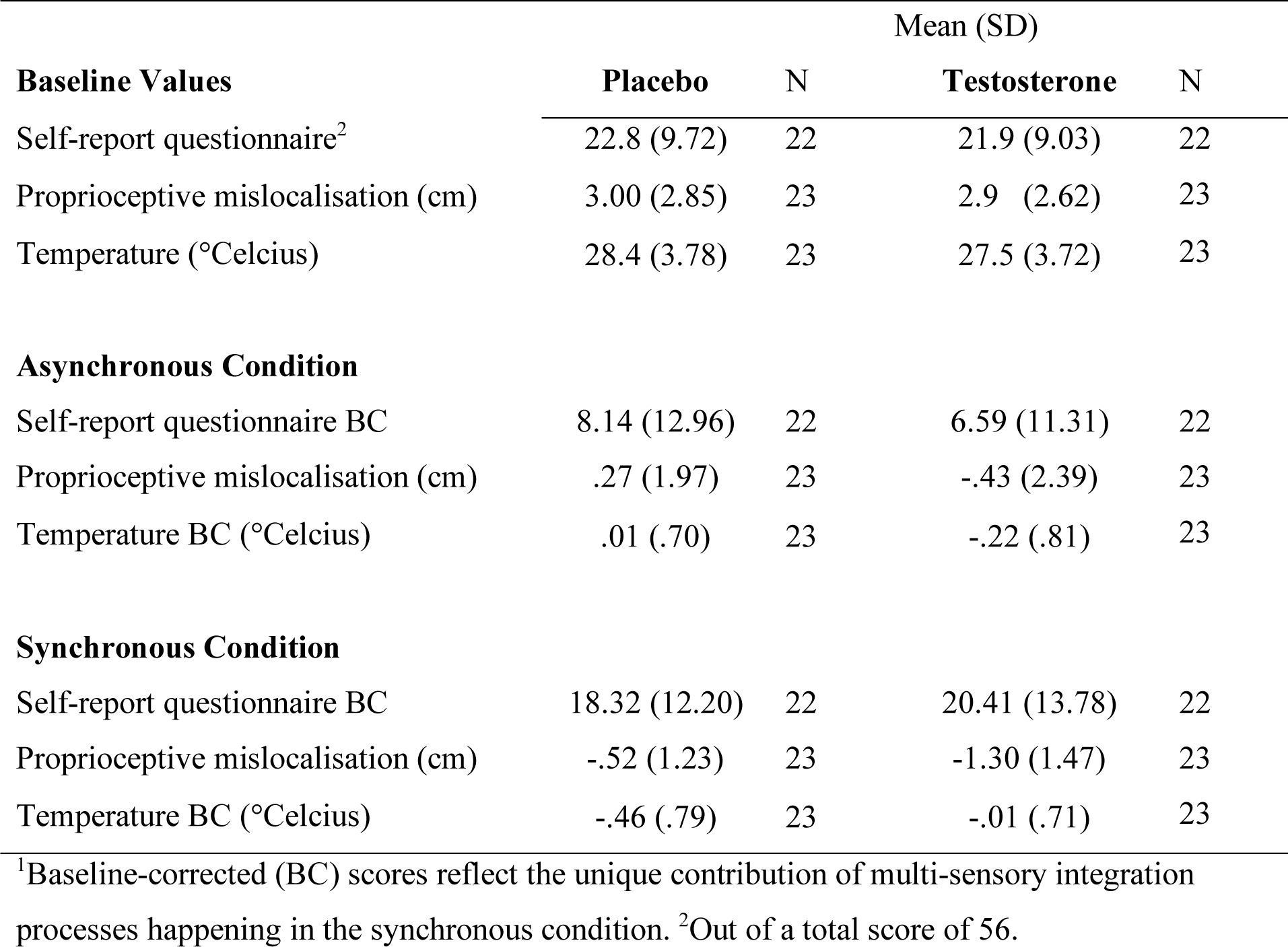
Baseline and baseline-corrected (BC) descriptive statistics for measures of the RHI

### 3.1 Self-report questionnaire

Mean scores on the self-report questionnaires were submitted to a mixed ANOVA, with stroking condition (visual-tactile stimulation) as the within-subjects factor and group (placebo versus testosterone) as the between-subjects factor. There was a significant main effect of visual-tactile stimulation (*F*_1,42_ = 38.111, *p<.*00). Across groups, scores for subjective experience of the illusion were higher following synchronous stroking, indicating that the procedure successfully induced the illusion. There was no main effect for group (*F*_1,42_ = .01, *p* = .93), nor was the interaction significant (*F*_1,42_ = .86, *p* = .36), suggesting that there was no specific effect of testosterone on either stroking condition. Looking at the self-report perceptual shift variables, an independent samples t-test showed no significant differences between the groups (*t*_42_ = -.94, *p* = .36, *d* = .28). These results indicate that testosterone had no effect on the subjective experience of the RHI.

We then looked to see if there was a difference between the two groups on the first five items of the self-report questionnaire, which focus more specifically on the subjective feeling of ownership over the rubber hand (the latter three items inquire about the subjective experience of limb location). Groups did not differ at baseline (*t*_42_ = .39, *p* = .69). Refer to Table 1 to view these items. ANOVA results indicated that there was a main effect of visual-tactile stimulation (*F*_1,42_ = 44.18, *p<.00*). Both placebo (*M*=14.18, *SD*=7.08) and testosterone (*M*=15.26, *SD*=8.30) groups showed substantially increased ratings of subjective ownership for the synchronous stroking condition. There was however no significant interaction effect (*F*_1,42_ = .70, *p*=.41) implying that neither group was specifically influenced by a multi-sensory effect. A two-tail t-test comparing the perceptual shift of these 5 subjective ownership items confirmed this finding, revealing no significant difference between testosterone and placebo groups (*t*_42_ = -.84, *p* = .41, *d* = .25).

### 3.2 Proprioceptive drifts

Average proprioceptive drifts were analysed using a mixed ANOVA, with stroking condition (visual-tactile stimulation) as the within-subjects factor and group (placebo versus testosterone) as the between-subjects factor. There was a main effect for stroking condition (*F*_1,44_ = 4.104, *p* = .04) and for group (*F*_1,44_ = 4.24, *p* =.04). Participants tended to perceive their hand as being closer to the rubber hand following synchronous stroking compared to asynchronous stimulation. Furthermore, collapsing across conditions, the testosterone group displayed significantly more proprioceptive drift than the placebo group. However, there was no interaction effect (*F*_1,44_ = .01, *p* = .94) and, confirming this, an independent samples t-tests of the proprioceptive drift perceptual shift variables showed no significant difference between the two groups (*t*_44_ = .08, *p* = .94, *d* = .02).

### 3.3 Temperature change

Scores reflecting the change in limb temperature were submitted to a mixed ANOVA, with stroking condition (visual-tactile stimulation) as the within-subjects factor and group (placebo versus testosterone) as the between-subjects factor. There were no main effects for stroking (*F*_1,44_ = 1.33, *p* = .25) or group (*F*_1,44_ = .32, *p* = .58); however, the interaction was significant (*F*_1,44_ = 9.020, *p* = .004). Independent samples t-tests (one-tailed) were used to compare the change in temperature between the two groups for each stroking condition. Following synchronous tactile stimulation, the temperature change between the testosterone and placebo groups was significant (t_44_ = −2.018, p = .02). In contrast, temperature change did not differ significantly between testosterone and placebo groups following the asynchronous condition (t_44_ = 1.03, p = .31). To further explore the interaction, paired sample t-tests (2-tailed) were used to test differences in temperature change scores within groups between stroking conditions. There was a significant difference between the synchronous and asynchronous stroking condition in the placebo group (*t*_22_ = −2.69, *p* = .01), but not in the testosterone group (*t*_22_ = 1.45, *p* = .16).

T-tests indicated that the drop in limb temperature from baseline to post-induction in the synchronous condition was significant in the placebo group only (*t*_22_ = 2.737, *p* = .006), who dropped by .457 degrees Celsius, but not in the testosterone group (*t*_22_ = .06, *p* = .43), where only a .009 decrease was observed. There were no significant differences between baseline and post-induction temperature scores in the asynchronous condition in the placebo (*t*_22_ = -.09, *p* = .93) nor the testosterone group (*t*_22_ = 1.29, *p* = .21).

Corroborating these findings, an independent samples, two-tailed t-test comparing the shift in temperature change from asynchronous stroking to synchronous stroking (temperature shift) showed a significant difference between testosterone and placebo groups (*t*_44_ = −3.003, *p =*.004, *d* = .89). These results indicate that limb temperature dropped significantly in response to synchronous tactile stimulation in the placebo group, but no such effect occurred after administration of testosterone.

Relationships between the perceptual shift indices of the RHI were assessed in the placebo group only. The self-report questionnaire was negatively correlated with temperature change (*r*(23) = .44, *p =* .04). However, after applying a Bonferonni correction for multiple comparisons, this finding was no longer significant. Looking specifically at the 5 items which focus on the subjective experience of ownership, there was a moderate to strong negative correlation with temperature drop, which remained significant after correcting for multiple comparisons (*r*(22) = -.57, *p =* .006). This finding implies that greater drops in limb temperature were associated with increased ratings of ownership over the rubber hand. Proprioceptive drift was not significantly related to subjective ownership (*r*(22) = -.30, *p =*.17) or temperature shift (*r*(23) = -.06, *p =* .78).

### 3.4 Mood data

There were no significant differences between the placebo and testosterone group on any subscale of the POMS, nor did the composite score, reflecting total mood disturbance (TMD), differ between the two groups (*t*_42_ = .29, *p* = .77). Correlation analyses of the mood data and indices of the RHI in the placebo group indicated that TMD was positively related to proprioceptive shifts toward the rubber hand (*r* (22) = -.59, *p = .*004). This result survived a Bonferroni correction which specifies an alpha of .016 for significance. TMD was not related to temperature changes (*r* (22) = .17, *p = .*44) nor subjective reports of ownership over the rubber limb (*r* (21) = .23, *p = .*32).

**Figure 3.**
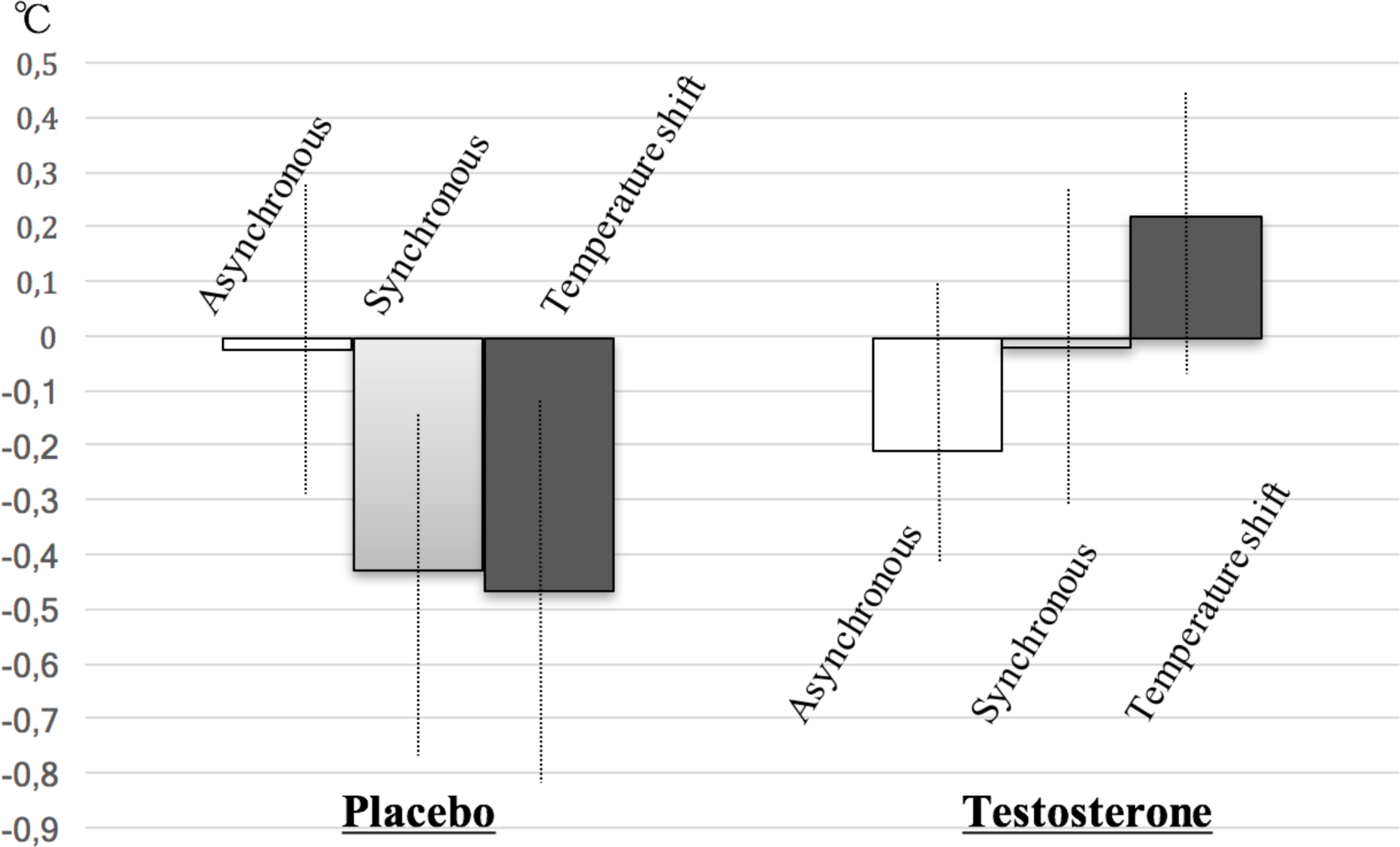
Effects of testosterone versus placebo on limb temperature change across conditions. During the Synchronous stroking condition, participants in the placebo group demonstrated a significant drop in limb temperature while no such drop occurred in the testosterone group. ‘Temperature shift’ is the difference between synchronous and asynchronous change and represents the unique contribution of multi-sensory processes. Error bars denote s.e.m.

## 4. DISCUSSION

Working within an embodied cognition framework, in this study we explored the hypothesis that effective maintenance of the body’s representational boundaries can be regarded, conceptually, as a defensive state facilitating the integrity of the self. We anticipated resistance toward the RHI in a sample of women administered 0.5mg of testosterone, a steroid hormone well known to promote defensiveness (Terburg et al., 2012; van der Westhuizen & Solms, 2015) and motivations for dominance and power (Carré & Putman, 2010; Eisenegger et al., 2011; van der Westhuizen & Solms, 2015a; Ronay & von Hippel, 2010). Testosterone had no effect on the subjective experience of the RHI or on measures of proprioceptive drift in comparison to placebo. However, the group of participants administered testosterone failed to show the same decline in limb temperature in response to synchronous tactile stimulation, as was seen in the placebo group. Thus, while explicit bodily representations were unaffected by testosterone, our hypothesis was confirmed in relation to implicit bodily representation, in that physiological processes related to thermoregulation operating outside of conscious awareness were maintained after administration of testosterone, but not after placebo.

### 4.1 Dissociable mechanisms

Consistent with several recent studies (Hohwy & Paton, 2010; Kammers, Rose & Haggard, 2011; Moseley et al., 2008; Thakkar, Nichols, McIntosh & Park, 2011; Tsakiris, Tajadura-Jiménez & Constantini, 2011; Van Stralen et al., 2014), induction of the RHI in the synchronous stroking condition lead to a significant decrease in limb temperature in the placebo group. The drop in temperature was of similar magnitude to that reported by Van Stralen et al. (2014) who employed an identical method. In their seminal study, Moseley et al. (2008) showed that cooling of the real hand is not a simple corollary effect of simultaneous visual-tactile input *per se*, nor due to a shift in attention away from the real hand, but that it is specifically the disruption in the subjective feeling of ownership that causes the drop in temperature. As such, they proposed that cooling should be interpreted as a disturbance in the physiological regulation of the real hand.

Not all studies report reliable or consistent changes in limb temperature (David et al., 2014; Dehaan et al. 2017; Grynberg & Pollatos, 2015; Paton et al., 2012; Rohde et al., 2013; Thakkar et al., 2011). Hohwy and Paton (2010) documented a drop in temperature from synchronous stroking, but not in all variants of the illusion that correspond with an equal strength in subjective ownership. Another study found a decrease in both the experimental and control conditions, leading these authors to argue that a thermal change is not a strict correlate of the experience of ownership (Rohde, Wold, Karnath & Ernst, 2013). Indeed, while the current findings show that reductions in temperature and feelings of ownership are significantly correlated under placebo, the finding that the cooling effect was not seen in the testosterone group supports the view that temperature changes and explicit feelings of ownership reflect dissociable mechanisms. Whether the two coincide may be determined by other factors.

It could be argued that implicit processes which directly correlate with bodily homeostasis have more immediate significance for the *re*-acting individual (eg. During social competition), unlike explicit awareness which arguably emerged later in phylogenetic development (Panksepp, 1998) and which is relatively superfluous to rapid motor-control (Cole & Paillard, 1995). This distinction may underlie a dissociation between homeostatic and subjective aspects of body ownership. Autonomic investment might be seen here as an indicator of active, first-person embodiment, that is, “I” (subject) versus “me” (object) (Babo-Rebelo, Richter & Tallon-Baudry, 2016). In support of this idea, Tsakiris Tajadura-Jiménez, and Costantini (2011) recently demonstrated that individuals with lower interoceptive awareness, a sensory channel argued to be critical in the foundation of stable bodily consciousness (Damasio, 2011), tend to react with larger drops in limb temperature in the RHI. This same group of individuals are known to experience body incompetence and the tendency to objectify the self (Ainley & Tsakiris, 2013). Our findings therefore suggest a role for testosterone in the maintenance of robust, *implicit* bodily representations that may support action. Explicit, body-as-object representations appear, on the other hand, to be less influenced by testosterone.

### 4.2 Testosterone and self-other boundaries

Several studies have interpreted findings from the RHI in terms of self-other discrimination, that is, the ability to distinguish “me” and “not-me” percepts, actions and feelings. Self-other blurring, especially at the level of external body representations, has been linked to conformity behaviour (Paladino et al., 2010) and the activity of oxytocin (Ide & Wada, 2017). It is possible therefore that maintenance of limb temperature in response to the RHI may function as a protective mechanism against self-other blurring. In support of this, in our placebo sample, we observed a significant negative association between subjective ownership and limb cooling, suggesting that subjective identification with a “not me” body part was associated with disinvestment of the bodily self.

However, the pattern of findings in the hormone-treated group does not suggest that testosterone promotes, necessarily, a *distinction* between self and other during induction of the illusion, since we report comparable scores of subjective feelings of ownership. Nor, unlike those observed in the placebo group, do the findings indicate self-other blurring, ie., when subjective identification with the rubber hand corresponds with homeostatic disembodiment of the real hand. Instead, our results in the testosterone group show a self-*self* pattern. Participants in this group demonstrated the same capacity to embody the rubber hand while maintaining physiological regulation of their own hand. It is as if this group “gained” a limb while for the placebo group, the real hand was replaced by the prosthesis.

Solely representing ‘self’ likely predisposes to egocentric ways of thinking. Previous work supports this view in that testosterone not only has detrimental effects on social cognition (Hermans et al., 2008; Montoya et al., 2013) but also on the ability to collaborate effectively with others in group settings (Wright et al., 2012). More interestingly, Overbeck and Droutman (2013) have argued that the use of the self as a heuristic for inferring the internal states of others is regularly seen in socially powerful people because of the strong link between the powerful and their roles in the leadership domain as the ones who are to represent the group at large. Being in power, or feeling powerful, may therefore prompt individuals to rely more on their own thoughts and sensations. This relationship may function in both directions. Thus, having a self-anchored frame of reference, which in the context of the RHI may reflect in the preservation of implicit body boundaries, appears to be closely linked to testosterone dynamics and positions of status in group hierarchies.

Despite findings being consistent with a self-self pattern in the hormone-treated group, it is nonetheless surprising that subjective ownership in the RHI was not decreased by testosterone since empathy has been established as a predictor of subjective ownership but tends to be hampered by the administration of testosterone (Hermans et al., 2008; Montoya et al., 2013). This calls into question a simple, straightforward link between self-reported feelings of ownership and empathy, suggesting that the role of embodiment in empathic processing is more nuanced. Perhaps it is only individuals who are able to maintain *some* level of self-differentiation that exhibit higher levels of social altruism. Indeed, the ability to differentiate between self and other is key for successfully empathising with other minds because a loss of self may lead to excessive negative affect and subsequently impoverished helping efforts (Decety & Sommerville, 2003).

### 4.3 Testosterone and the preservation of the bodily self

From a Bayesian brain perspective (Pezzulo, Rigoli & Friston, 2015), top-down models that guide action must be anchored to the bodily, interoceptive self if they are to succeed in serving power and self-interests. Without doing so, predictive models of the self would be ineffective. Maintaining thermal homeostasis in the context of the RHI may help therefore to ensure stability and unity of the agentive, bodily self via enabling access to “gut feelings” (Dunn et al., 2010).The putative role for testosterone in the preservation of the bodily self is consistent with numerous studies associating the hormone to defensive, self-serving behaviour in humans and animals (Eisenegger et al., 2011; Dreher et al., 2016; Terburg et al., 2012; Wright et al., 2012). Via its influence on the amygdala and insular cortex, for instance, testosterone sustains threat monitoring (Bos et al., 2010; Hermans, Ramsey & van Honk, 2008). Since the insula has also been linked to sensory-motor agency (Farrer & Frish, 2002) and body ownership (Karnath & Baier, 2010), the effect of testosterone on homeostatic resistance to RHI manipulations may be mediated, in part, by insula activity. This link reinforces the proposal that defence of body boundaries might be recruited in the service of more complex forms of motivated, defensive behaviour.

Of the studies performed on the RHI to date, none have directly explored the emotional correlates of temperature changes. It is therefore difficult to align the current findings of testosterone with prior research in the field. However, in other contexts, there appears to be a reliable link between reductions in skin temperature of the limbs and negative, largely disempowering emotional states. Several studies report drops in surface skin temperature in response to cognitive or emotional stress (Boudewyns, 1976; Bugental & Cortex, 1998; Mittleman & Wolff, 1939; Or & Duffy, 2007; Zhai & Barreto, 2006) and music that elicits negative emotional feelings (McFarland, 1985). Rimm-Kaufman and Kagan (1996) report a decrease in limb temperature in response to threatening personal questions but not to cognitive tasks or fear-eliciting film clips. Corroborating this, social ostracism has also been found to lower skin temperatures (Ijzerman et al., 2012), suggesting perhaps that a cooling of the body contributes to the implicit understanding of what it means, biologically and psychologically, to be socially threatened and disempowered. This link may apply exclusively to effector limbs, since increases in cheek temperature, as in the case of blushing, tends to be associated with social anxiety (Voncken, & Bögels, 2009; see also Nakanishi & Imai-Matsumura, 2008). Indeed, within the body, a drop in skin temperature appears to be related to reduced blood flow as a result of vasoconstriction which occurs following activation of the sympathetic nervous system (Wallin, 1981), i.e, the central stress response. It is possible, that on an implicit level, the RHI manipulation is experienced as a stressor, given that it threatens bodily integrity, and in this way may activate the stress response. Most notably, testosterone administration has been shown to attenuate responsiveness of the integrated stress system (Hermans et al., 2007), possibly explaining its ability to maintain skin temperature. Plausibly, such thermoregulatory effects would impact upon the sensation of effort, which is a key phenomenological aspect of motor control (Pacherie, 2008). Together, these findings imply that maintenance of body temperature is psychologically stabilising. The advantages to social coping achieved by those with higher levels of testosterone (Edwards, Wetzel & Wyner, 2006; Enter, Spinhoven & Roelofs, 2014; Rowe et al., 2004; van Honk et al., 2004) may thus derive from similar embodied mechanisms.

Finally, the effect of testosterone on preserving bodily representations during the illusion is also consistent with the finding that larger errors in proprioception, as a result of multisensory integration, were related to significantly higher levels of mood disturbance in the placebo group. Even though we did not observe any influence of testosterone on mood or proprioception directly, these findings nonetheless indicate that emotional factors do impinge on bodily representations. Specifically, here we show that negative mental states appear to “loosen” the bodily schema. Similarly, Kállai et al. (2015) report an association between *harm avoidance* and proprioceptive drift. That we did not observe any relationship between temperature shift and mood, or between testosterone and proprioceptive mislocation, suggests that thermoregulatory changes in the context of the RHI may reflect unconscious or implicit mental states. For instance, testosterone in particular is known to produce many of its emotion-regulating functions outside of conscious awareness (van Honk, Peper & Schutter, 2005). This might explain why it has no specific effect on proprioception during multisensory integration. It is, however, possible that the relationship reported here between mood items of the POMS and proprioceptive drift simply reflect reduced attention as a result of the negative mood states. As such, future studies will benefit from incorporating affective variables in RHI studies to advance our understanding of the emotional mechanisms influencing embodiment.

### 4.4 Limitations

The current findings should be considered with the following limitations in mind. Although in keeping with many other testosterone administration studies (Enter et al., 2014; Terburg et al., 2012; van Honk et al., 2011), the sample size used here is relatively small. A larger sample, or a within-subjects design, may yield enough power to reveal more persuasive results, especially with regard to correlation analyses. Secondly, it is customary to assess circulating levels of testosterone, either in saliva or blood, to confirm significantly elevated levels of testosterone in the hormone treated group. We were not able to assess basal testosterone. However, based on previous studies that demonstrate a ten-fold increase in women’s total testosterone in response to 0.5mg of the hormone (Tuiten et al., 2000), we can infer with a reasonable degree of confidence that the significant difference in limb temperature between the hormone and placebo treated group during the RHI reflects activity of testosterone.

Though the current findings cannot determine conclusively whether the maintenance of limb temperature derives from peripheral/bodily or central/brain mechanisms, it is worth noting that testosterone plays a key role in influencing the body by increasing muscular and skeletal strength (Bhasin et al., 1996) and by increasing the excitability of motor neurons (Maeda & Pascual-Leone, 2003; Ziemann, 2004). These metabolic and neural processes could be argued to influence thermoregulation in general. However, here we only observed a specific effect of testosterone on temperature in response to synchronous stroking. No such effect emerged in the control condition, implying that the effect of testosterone on thermoregulation is probably explained instead by a change higher up in the nervous system. In support of this view, testosterone has been found to restore innervation of the peptide, arginine vasopressin, in the nucleus of the solitary tract (NST; Goudsmit, Fliers & Swaab, 1988), a key structure of the brain’s interoceptive system receiving afferent input from the body periphery (Craig, 2003). The NST shows considerable androgenic immunoreactivity (Hamson, Jones & Watson, 2004) and furthermore, the brain’s core temperature regulator – the medial pre-optic nucleus of the hypothalamus – receives projections directly from the NST (Ricardo & Koh, 1978). It therefore seems reasonable to assume that the preservation of thermoregulation by testosterone in the experimental limb derives from activity in the brain.

### 4.5 Concluding remarks

In our sample, participants administered 0.5mg of testosterone failed to demonstrate the cooling of their own hand in response to the RHI, as was documented in the placebo group and several other recent studies (Hohwy & Paton, 2010; Kammers, Rose & Haggard, 2011; Moseley et al., 2008; Thakkar, Nichols, McIntosh & Park, 2011; Tsakiris, Tajadura-Jiménez & Constantini, 2011; Van Stralen et al., 2014). Previous authors have proposed that cooling should be understood as a disturbance in physiological homeostasis. Drawing on embodied cognition theory, we propose that thermal regulation contributes to the effective maintenance of the body’s representational boundaries and can be likened to a defensive state facilitating the unity of the self. Certainly, if not in reference to the self (i.e. without interoceptive investment in one’s own body), the empowering nature of embodiment simply cannot occur. Indeed, one’s own body is a space-occupying entity with an “inside” and an “outside” and, like any other territory, if the integrity of these boundaries are efficiently monitored, it provides phenomenological stability and a clear boundary for monitoring external threat. More significantly, this provides a means via which to expropriate valuable objects from the outside world via a process of self-association (De Dreu & Knippenberg, 2005; Turk et al., 2011). Based on our finding that testosterone prevents thermal deregulation during the RHI, we therefore speculate that the mechanism regulating homeostatic control over the body during multi-sensory integration in the RHI is an emotional one linked to self-preservation.

## 5 Acknowledgements

DvdW received support from the Oppenheimer Memorial Trust and South Africa’s National Institute for the Humanities and the Social Sciences.

